# Mechanisms of strain diversity of disease-associated in-register parallel β-sheet amyloids and implications about prion strains

**DOI:** 10.1101/472712

**Authors:** Yuzuru Taguchi, Hiroki Otaki, Noriyuki Nishida

## Abstract

The mechanism of strain diversity of prions still remains unsolved, because the investigation of inheritance and diversification of the protein-based pathogenic information demands identification of the detailed structures of abnormal isoform of prion protein (PrP^Sc^), while it is difficult to purify for analysis without affecting the infectious nature. On the other hand, the similar prion-like properties are recognized also in other disease-associated in-register parallel β-sheet amyloids including Tau and α-synuclein (αSyn) amyloids. Investigations into structures of those amyloids by solid-state nuclear magnetic resonance spectroscopy and cryo-electron microscopy recently made remarkable advances, because of their relatively small sizes and lack of post-translational modifications. We review the advances on those pathogenic amyloids, particularly Tau and αSyn, and discuss their implications about strain diversity mechanisms of prion/PrP^Sc^ from the viewpoint that PrP^Sc^ is an in-register parallel β-sheet amyloid. We also present our recent data of molecular dynamics simulations of αSyn amyloid, which suggest significance of compatibility between β-sheet propensities of the substrate and local structures of the template for stability of the amyloid structures. Detailed structures of the αSyn and Tau amyloids are good surrogate models of pathogenic amyloids including PrP^Sc^ to elucidate not only the strain diversity but also their pathogenic mechanisms.

---

Strain diversity is one of the most mysterious feature of prions. The strain-specific traits of prions are enciphered in the structures of the abnormal isoform prion protein (PrP^Sc^), and they are stably inherited over generations solely by passaging the exact structures of the PrP^Sc^ through template-directed refolding of the normal isoform prion protein (PrP^Sc^), where the template PrP^Sc^ imprint the structural details onto the substrate PrP^C^. Moreover, the strain-specific pathogenic information encoded in the conformation of the PrP^Sc^ is reproducibly “translated” into the strain-specific clinicopathological traits in the manifested diseases [1][2]. This view is widely accepted as the protein-only hypothesis but detailed mechanisms, e.g., specifically what structures of PrP^Sc^ encodes the pathogenic information and how the information is translated, remain a challenge to the current biology. Investigations of the storage, inheritance and diversification of the protein-based pathogenic information demand identification of the structure-phenotype correlations, but it is very difficult because detailed structures of the entire PrP^Sc^ is not available yet due to its incompatibility with high-resolution structural analyses, i.e., X-ray crystallography or nuclear magnetic resonance spectrometry (NMR). The incompatibility is attributable to difficulty in purification without losing infectivity [3], difficulty in recapitulating bona fide prion in vitro with infectivity and toxicity, and relatively large sizes of PrP with post-translational modifications. Solid-state NMR (ssNMR) revealed in-register parallel β-sheet structures of in vitro-formed amyloids of the peptide corresponding to the residues 23-144 [4], but the structures of the whole molecule would be necessary for elucidation of the mechanism of the strain diversity. On the other hand, cryo-electron microscopy (cryo-EM) of PrP^Sc^ lead to the four-rung β-solenoid model [5], although the resolutions were not high enough for atomic-level modeling. Which model is more plausible is still controversial. As for the translation of the strain-specific structures, interactions of PrP^Sc^ with environments and/or other proteins are essential because each strain recognizes the preferable cell groups to manifest the strain-specific lesion profiles, which provide the environments and factors required for efficient propagation [6][7][8].

Many clinically important neurodegenerative diseases are caused by disease-associated amyloids, e.g., Alzheimer’s disease (AD) by β-amyloid, Parkinson’s disease (PD) and dementia with Lewy bodies (DLB) by α-synuclein amyloids, and tauopathies including Pick’s disease by Tau amyloids. Interestingly, they show “prion-like properties” including transmissibility to animals and even strain diversity [9][10][11]. Although they have no homology in the amino acid sequences, those amyloids share a common basic architecture, in-register parallel β-sheet structure [12][10][13], which is characterized by a stack of β-loop-β motifs with the identical types of amino acids aligned along the fibril axis with narrow intervals, 4.8 Å (**Fig 1A**). A given residue of the amyloid therefore can interact most frequently with the identical-type counterparts of the adjacent layers. This structural feature may accentuate the characteristics of each residue of the peptide and greatly contribute to determination of the amyloid conformations, e.g., hydrophobic residues make extensive hydrophobic patches spanning along the entire fibril axis, while charged residues may disorder the local structures by repulsion between the layers and be preferentially exposed to the solvent. The feature also explains why a single mutation greatly affects properties of the amyloid because it replaces the entire column (**Fig 1B**). Why do different proteins take the same in-register parallel β-sheet structures that can efficiently propagate? The in-register amyloids are hypothesized to have lower free energy than the native conformations [14]. The hydrogen bonds between the backbones of the amyloid thermodynamically favor the in-register alignment [15]. Roterman et al. regarded the in-register amyloid as a “ribbon-like micelle” whose exposed hydrophobicity at the stack ends enable endless elongation [16]. Efficient propagation of the amyloids may stem from those thermodynamic features of the in-register parallel β-sheet structures. If the structures of the amyloid are autonomously determined depending on the primary structure and thermodynamic principles, in-register parallel β-sheet amyloids are suitable experimental objects for MD simulation.

**Fig 1.**
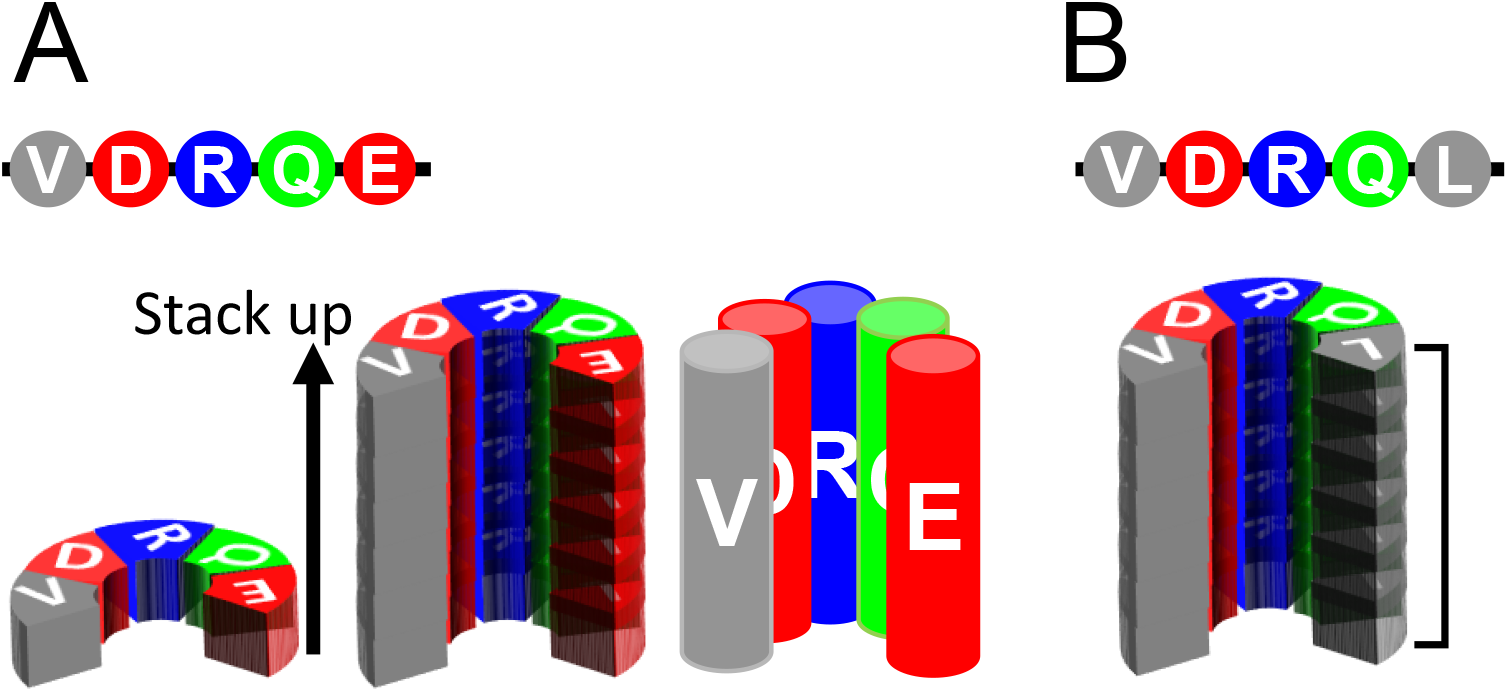
Unique structures of in-register parallel β-sheet structures. **A**. Schematics illustrating the primary structure of a peptide (top), a layer of the peptide in β-sheet structure (lower left), a stack of the in-register parallel β-sheets of the peptide (middle). Note that the identical types of amino acids pile up along the fibril axis of the amyloid with intervals ∼4.8 Å, and they can be compared to columns of the same types of amino acids (right panel). **B**. An example of a substitution mutation of the peptide, Leu for Glu (top). The substitution replaces a whole column (bracket) from the negatively-charged to the hydrophobic, creating a large hydrophobic patch encompassing the length of the fibril.

Recent progresses in the structural analyses of the disease-associated amyloids are even greater than those of PrP prions, partly because of their relatively small sizes, lack of post-translational modifications, and in-vitro reproducibility of the cytotoxicity and infectivity of the amyloids [17]. Resolutions of the structures of in-register amyloids determined by ssNMR or cryo-EM [13][18][19][20][10] are high enough to allow analyses of structure-phenotype relations of those amyloids. Here, we review those recent advances in investigations of the structures and strain diversity of disease-associated in-register parallel β-sheet amyloids, introduce our in-silico data on αSyn amyloid, and discuss whether the knowledges from those amyloids are applicable to PrP^Sc^. For the purpose, we particularly focus on relatively large pathogenic amyloids, α-synuclein and Tau.

## Progresses in investigation of Tau amyloids

Tau protein is abundantly expressed in the nervous systems, especially in nerve cell axons, and intrinsically disordered with a low content of secondary structures [21]. Tau has six isoforms depending on patterns of alternative splicing of the exons 2, 3, and 10. The exon 10 corresponds to the second repeat of the four-repeat motifs of the C-terminal microtubule-binding domain, and the absence or presence of the exon 10 results in “three-repeat” (3R) or “four-repeat” (4R) Tau, respectively. The repeat motifs converts to an in-register parallel β-sheet structures in tauopathies. Interestingly, which of the six isoforms preferentially convert to amyloids varies depending on disease types: for example, in Pick’s disease the amyloids consist predominantly of the 3R-Tau, whereas AD amyloids contain both 3R- and 4R-Tau [22][23].

A recent great progress regarding the mechanism of strain diversity of amyloids was the identification of detailed structures of two different Tau amyloids isolated from diseased brains of AD and Pick’s disease by cryo-EM [10][20]. The distinct structures of those Tau amyloids demonstrated unequivocal examples of strain-specific structures. One of the notable differences between AD Tau and Pick’s Tau amyloids is the positions of acute-angle β-arches, which reminisces the mechanism of strain diversity of a yeast prion Sup35 postulated by Kajava et al. [24]. Interestingly, except one β-sheet which is straight in the Pick’s fold but bent in the AD fold making a new β-arch, many of the β-sheet regions in the AD amyloid maintained the β-sheet structure also in the Pick’s amyloid as if the positions of β-sheets are fixated, albeit different facing partners for cross-β spine formation. The primary structure might be significant determinants of positions of β-sheets in in-register parallel amyloids, and cross-β-spine formation would be necessary to stabilize the β-sheets by concealing hydrophobic residues from the water [25]. Consistent with the different patterns of the cross-β spines, Pick’s form and the AD form of Tau amyloids showed different proteolytic-fragment patterns on immunoblots after digestion with trypsin [26].

Falcon and colleagues suggested mechanisms of the strain diversity of Tau amyloids which is applicable to other amyloids as well. They postulated that steric conflicts of the branching Cβ of Val_30_0 of 4R-Tau with the omega-like structure formed by Pro_2_7_0_-Gly-Gly-Gly_2_7_3_ of the 3R-Tau of Pick’s amyloid make the 4R-Tau an incompatible substrate, hampering cross-seeding of 4R-Tau by the Pick’s amyloid [22]. This type of strain barrier due to incompatibility of local structures between the substrate monomer and the template amyloid is discussed below again regarding αSyn and PrP^Sc^.

Shifts of pairs of the cross-β spines accompanied by alterations in the positions of β-arches enable generation of structural polymorphs even by amyloids with relatively simple conformations like Tau amyloids. The similar mechanisms therefore can be operative in diversification of strains of other amyloids including PrP^Sc^. For example, it could be one possible explanation for the mechanism of how smaller proteolytic fragments of 12 or 13 kDa of PrP^Sc^ are produced depending on the disease types [27][28][29]. If the fragments are produced by cleavage at relatively protease-sensitive sites of PrP^Sc^, their differential sizes imply a shift of the protease-sensitive sites and such a shift can occur by alterations of cross-β spine patterns and β-arches. For example, when conversion to the β-sheets and formation of cross-β spines propagate to the whole molecule like a slider and zipper from an amyloid core/interface that is first converted by the template amyloid, it is conceivable that the position of the amyloid core/interface determines the pairing partners of the cross-β spines, and a process starting from another amyloid core results in a distinct pattern of cross-β spines. Indeed, infectious PrP^Sc^ can be induced by either type of in vitro-formed PrP fibril with protease-resistant core in the N-terminal [30] or C-terminal region [31]. The Creutzfeldt-Jakob diseases (CJD) with 13-kDa fragments, e.g. sporadic CJD with methionine homozygosity at the codon 129, showed shortened incubation periods in human-mouse PrP-expressing mice, corroborating the significance of the structural variation in strain barriers of prion [29][32].

## Progresses in investigation of αSyn amyloids

αSyn is a cytosolic protein abundantly expressed in neurons, representing ∼1 % of the total cytoplasmic proteins [33]. Since αSyn was cloned as the “non-Aβ component” (NAC) from the AD amyloids [34], the first-identified region encompassing residues 61-95 is often referred to as the NAC region. It is localized mainly in presynaptic nerve terminals and contributes to controls of the neurotransmitter releases [35]. In addition to the diversity of clinicopathological pictures among synucleinopathies, i.e., PD, DLB, and multiple system atrophy (MSA), familial PD show clinical variations depending on the causative mutations in the SNCA gene, regarding age of onset, disease duration, and presence of pyramidal signs [36][37]. Variations in physicochemical properties of αSyn amyloids are also well-documented. For instance, in-vitro formed αSyn fibrils show at least two types of morphologically-distinguishable strains, “ribbon” and “fibril” types, which are different in dimensions of the fibrils, blotting patterns of protease-resistant fragments, cytotoxicity, and optimal salt conditions for efficient in-vitro propagation [38][39]. Their secondary structures determined by ssNMR revealed strain-specific β-sheet distributions: the ribbon had stable β-sheet structures in the N-terminal region encompassing the residues 1 to 38, whereas the corresponding region of the fibril type was disordered except for a short β-sheet 16-20. To the contrary, the residues 44-57 was disordered in the ribbon while they are in β-sheet in the fibril type. Although the distal NAC region of the fibril type might be structurally varied [39], positions of β-sheets in the NAC regions were relatively similar between the ribbon- and the fibril-types [40][41], reminiscent of Pick’s and AD’s Tau amyloids sharing many β-sheet regions. Besides in vitro-formed fibrils, αSyn amyloids derived from DLB- and MSA-affected brains also differed in the blotting patterns of proteolytic fragments [42]. Like the different optimal propagation conditions between the ribbon and the fibril, MSA-αSyn amyloid is formed in the milieu of oligodendrocytes, whereas DLB-αSyn amyloid is formed in the neurons [42].Thus, αSyn amyloids can be a good model in investigation of strain diversification of pathogenic amyloids because of those unequivocal structural differences between strains and the structure-phenotype correlations as suggested by the proteolytic-fragment patterns.

Another vigorously attempted biochemical approach for structural information of αSyn amyloids is assessment of cross-seeding efficiencies between αSyn molecules with familial-PD-associated mutations, e.g., between A30P and A53T [43]. Evaluation of aggregation formation and cross-reactions of in vitro-translated GFP-fusion mutant αSyn demonstrated that αSyn mutants with A53T, H50Q, or E46K spontaneously form large aggregates and efficiently co-aggregate with each other, whereas they do not co-aggregate with the wild-type or mutants with G51D or A30P; the latter group of three αSyn species formed smaller aggregates and co-aggregate each other within the group [44]. Those findings suggested existence of two mutually exclusive aggregation paths of αSyn amyloid formation. Cross-seeding experiments also proved the contribution of the N-terminal- and the C-terminal-side regions to efficient propagation and strain-specific properties of αSyn amyloids, as demonstrated by in vitro fibril formation and in vivo inoculation experiments with various human-mouse chimeric αSyn [45], or with N- and C-terminally truncated mutant αSyn [46]. Not only cross-seeding efficiencies between αSyn molecules, strain-specific structures of αSyn amyloids can also affect interactions with other proteins. For instance, induction of Tau pathologies by αSyn preformed fibrils is reported to be a strain-specific phenomenon. In vitro-formed two different strains of αSyn amyloids with distinct proteolytic-fragment patterns exhibited different efficiencies in cross-seeding Tau and induction of phosphorylated Tau in vivo, and the N-terminal region was important for the cross-seeding of Tau [47]. Whether cross-seeding of other proteins by αSyn amyloids, e.g., Aβ [48] or amylin [49], are also strain-dependent would be also intriguing.

Atomic-level structures of in vitro-formed αSyn amyloids were determined either by ssNMR or cryo-EM [50][19][18]. The conformation determined by ssNMR exhibited a “Greek-key” conformation encompassing the residues 35 to 99 with intricate interactions among the constituent β-sheets [50]. Although the majority of αSyn fibril observed in PD brains are intertwined two-protofibril forms [51], the Greek-key αSyn amyloid was rather stable in MD simulations of 400 ns even without another protofibril (**Fig 2A and 2B, middle and right panels**) [52]. Regardless of relative instability in the region 47-61 of the stack-end molecules, the overall stability of the amyloid stack in the conformation was sufficient for evaluation of influences of various mutations, e.g., G51D or A53T, on the amyloid structures [52]. The detailed structures of αSyn amyloids determined by cryo-EM revealed co-existence of two apparently distinct types of fibrils formed under the same conditions. Both the polymorphs, “rod” and “twister”, consisted of inter-twined two protofibrils and shared the common amyloid “kernel”, but they had different inter-protofibril interfaces [19]. The authors therefore postulated that the inter-protofibrilar interface might be an important determinant of αSyn amyloid strains. Although the rod type and the one reported by Guerrero-Ferreira et al. (PDB ID: 6h6b) had Greek-key conformations, they were slightly different from the one identified with ssNMR (PDB ID: 2n0a) in that the Ala53 pointed outward to constitute the inter-protofibril interface and that the highly-charged region 56-60 formed a decent β-sheet. Whether those two polymorphs represent two different strains, rather than structural polymorphs of the same strain with the same kernel, would need to be corroborated by strain-specific biological properties of each polymorph.

**Fig 2.**
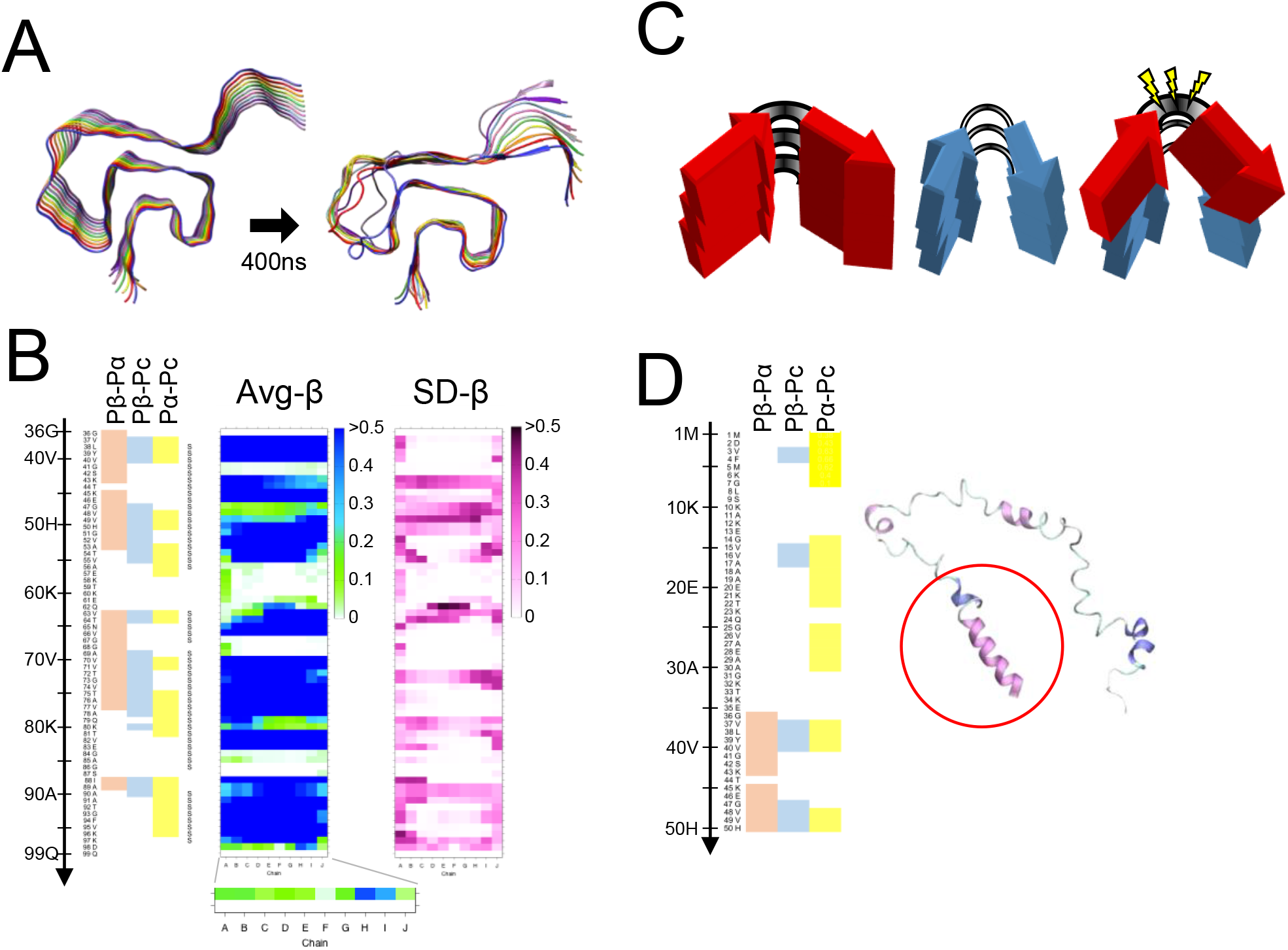
Correlation between the results of MD simulation and of the secondary-structure prediction. **A**. The “Greek-key” conformation of αSyn amyloid (PDB ID: 2n0a) (left). The status of the stack of αSyn amyloid after 400 ns of the MD simulation (right). The protocol for the MD simulations are the same as we previously used (Ref. 52) **B**. (Left) Propensity profiles predicted from the primary structure of αSyn by a neural network secondary structure prediction (Ref. 53). The heatmap exhibits the magnitude correlations between the conventional β-sheet propensity (Pβ), α-helix propensity (Pα) and coil propensity (Pc) by the new set of parameters, (Pβ-Pα), (Pβ-Pc) and (Pα-Pc). Zero- and positive-value residues are indicated with the color codes, red, blue and yellow, respectively. (Middle) A heatmap of the average β-sheet propensity of the αSyn amyloid observed in the MD simulation of five independent runs. The color codes indicates the proportion of time when a given residue is in β-sheet structures during the simulation run. For all the heatmaps of average β-sheet propensity and SD, the vertical and horizontal axes represent the residues from 36-99 and the ten layers/chains (A-J) of the amyloid stack, respectively. (Right) A heatmap of the standard deviation (SD) values of β-sheet propensities of the αSyn amyloid observed in the MD simulation of five independent runs. **C**. A schematic illustrating a hypothetical local incompatibility in cross-seeding between two heterologous amyloidogenic peptides. The red peptide has a non-flexible loop region, whereas the blue peptide has a more flexible loop. When the red peptide with rigid loop is cross-seeded by the blue-peptide amyloid, the substrate red peptide might undergo strains from the incompatibility between the intrinsic properties and the actual structures. **D**. A final snap shot of native-form αSyn (PDB ID: 2kkw) after 5 ns of MD simulation without any micelle: Only the N-terminal side encompassing 1-100 was used. Note that the N-terminal region with the high (Pα-Pc) values relatively maintain the α-helix (red circle).

## Insights from MD simulations of αSyn amyloids: significance of the intrinsic propensities and the local structures

The detailed structures of Tau and αSyn amyloids suggested strain diversification by alterations in positions of β-arches, pairing patterns of cross-β spines and inter-protofibrilar interfaces, which greatly depend on intricate interactions among the constituent β-sheets. On the other hand, the incompatibility between Pick’s Tau amyloid and 4R-Tau substrate by steric conflicts by the Cβ carbon of Val_30_0 demonstrated that strain barriers can be also posed by such local factors. We had hypothesized that conflicts between the local structures of the template amyloid and the corresponding local intrinsic propensity of the substrate peptide can cause a strain barrier, and that the intrinsic propensities are predictable by a secondary structure prediction algorithm [53][54]. Although the algorithm was originally designed for monomeric proteins, the singular structural feature of in-register parallel β-sheet amyloids could allow the application (**Fig 1A**). By comparing the predicted propensities, i.e., β-sheet propensity (Pβ), α-helix propensity (Pα) and coil propensity (Pc), of PrPs from various species, we hypothesized that compatibility of β-arches between PrP^Sc^ and PrP^C^ contributes to species barriers (**Fig 2C**) [54]. Here we test the hypothesis the local conformations and the propensity of substrate with MD simulation, which enables observation of the influences exclusively of the conformations on behaviors of the peptides without any any interference from other factors or proteins.

We chose the “Greek-key” αSyn amyloid (PDB ID: 2n0a) [50] as a representative model of an in-register parallel β-sheet amyloid, because of its intricately-interacting β-sheets and paucity of aromatic amino acids whose pi-pi interactions are not fully represented by the used force field. First, we compared predicted propensities with the results of MD simulations of the wild-type (WT) αSyn amyloid [52], to assess relations of the predicted propensities and the actual structures of αSyn amyloids in silico. Here we used a set of modified parameters, Pβ-Pα, Pβ-Pc and Pα-Pc, to clarify the magnitude correlations between the conventional parameters. These predicted propensities exhibited some correlation with the actual average β-sheet propensity from the MD simulations (**Fig 2B**), suggesting certain influences of intrinsic propensities on structures of the amyloid. Positive (Pα-Pc) values (**Fig 2B**, yellow areas) indicate high α-helix propensities in theory, and indeed the N-terminal region with positive (Pα-Pc) values maintained a helix in the MD simulation of a native αSyn (PDB ID: 2kkw [55]) even in the absence of micelles (**Fig 2D**). The β-sheet in the region 89-95 with positive-(Pα-Pc) values (**Fig 2B**) reminisces contribution of high α-helix propensities to amyloid formation which explains why amyloidogenic proteins often have α-helices in the native conformations, e.g., Aβ and PrP [56][57][58][59]. Given the correlation of the predicted propensity profiles with the structures of αSyn amyloid in silico, we next introduced substation mutations of isoleucine around the three loops encompassing the residues 56-62 [loop(56-62)], 67-68 [loop(67-68)] and 84-87 [loop(84-87)] (**Fig 2A**), specifically Glu61Ile (E61l), Asn65Ile (N65I), and Gly84Ile (G84I), respectively (**Fig 3A**). Then, we analyzed their predicted propensities. G61I locally raised (Pβ-Pα) and (Pβ-Pc) values (**Fig 3A**, middle). N65I raised (Pβ-Pc) in the loop(67-68) and the adjacent β-strands (**Fig 3A**, right). G84I raised (Pα-Pc) to positivity through the loop and changed the positive-(Pβ-Pc) spot at the residue 80 to the wider one encompassing 82-84 (**Fig 3B**).

**Fig 3.**
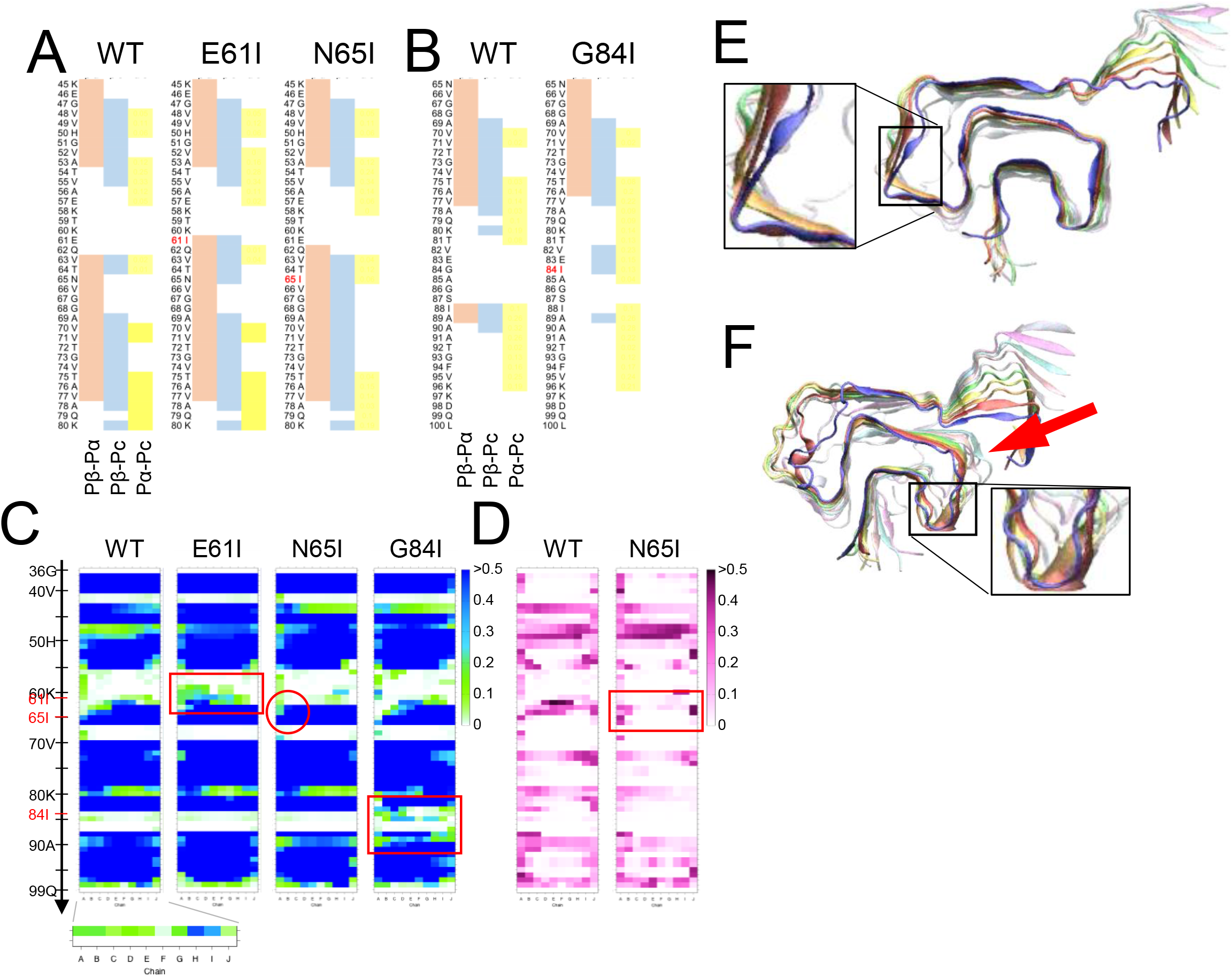
Inappropriately high β-sheet propensity in the loop region can destabilize the αSyn amyloid. **A & B**. Comparison of the predicted propensity profiles between WT-, E61I-, N65I- and G84I-αSyn. The mutation sites are indicated in red letters. Isoleucine substitutions were introduced near the loop regions based on the assumption that loops with higher β-sheet propensities may be less flexible. **C**. Comparison of heatmaps of average β-sheet propensities between WT-, E61I-, N65I- and G84I-αSyn. They represent five (for WT) or three (for the others) independent simulations. Note that all the mutants have higher average β-sheet propensities in the regions comprising the respective mutations (more green/blue cells in the red boxes for E61I and G84I, and in the red circle for N65I). **D**. Comparison of the heatmaps of SD values between WT- and N65I-αSyn. They represent five and three independent runs, respectively. Note that the region comprising the N65I mutation shows more stable β-sheet with smaller SD than WT-αSyn (red box). **E**. A final snap shot of E61I-αSyn after 400 ns of simulation. Note that β-sheets are newly induced in the loop region encompassing 56-61 (inset). The mutation tended to stabilize the global structures of the amyloid and occasionally induced such β-sheets in the loop. **F**. A final snap shot of G84I-αSyn after 400 ns of simulation. The mutation destabilized the structures around the mutation and also the adjacent regions (arrow). Occasionally β-strands were temporarily induced in the loop (inset). The differential effects of the isoleucine substitutions on the amyloid structures might be attributable to the shapes of the loops. The loop(56-62) is long and makes obtuse angles with the flanking β-strands. The loop(67-68) is short but rather flexible with two glycine residues in a row. The loop(84-87) is a U-shaped loop with a relatively-small turning radius, which would not accommodate newly-induced β-sheets.

MD simulations of homo-oligomers of those mutant αSyn were performed with the same conditions as we previously did [52], influences of N65I were subtle but the β-sheet encompassing 62-66 was more stabilized particularly on the chain-A side (**Fig 3C**, red circle) with smaller SD values than those of WT (**Fig 3D**); E61I occasionally induced β-strands in the loop(56-62) and tended to stabilize the amyloid stack (**Fig 3E**, inset). In contrast, G84I substantially destabilized the loop(84-87) and the adjacent structures (**Fig 3F**, arrow). In accordance with the raised local β-sheet propensity, β-strands were temporarily induced in loop(84-87) (**Fig 3F**, inset) but could not form a stable β-sheet. Those varied effects of isoleucine substitutions suggest that raised β-sheet propensity either stabilizes or destabilizes the loops depending on their shapes, lengths and flexibility: a “U-shaped” loop with small turning radius like loop(84-87) might be destabilized because the loop is too small to accommodate the β-sheet newly induced by the high β-sheet propensity. This provides a proof-of-concept to the hypothesis that incompatibility between the intrinsic propensities of the substrate with the actual local structures of the template amyloid affects stability of the amyloid [54]. To note, the amyloids contained only mutant αSyn without any mismatch in the primary structures among the peptides in the stack. In seeding reactions of amyloids or prions, homology in the primary structures between the seed and the substrate is an important determinant of seeding efficiency, but the present result corroborated that compatibility of intrinsic propensity of the substrate with local structures of the amyloid is more important than the sequence homology. This notion helps interpret the singular inter-species transmissions of prions where even the secondary transmissions are inefficient despite the homology between the PrP^Sc^ in the inoculum and the host PrP [60][61]. Interestingly, in contrast to the stabilizing effects of E61I or N65I in the home-oligomers, they showed destabilizing effects in hetero-oligomers combined with the wild-type αSyn (data not shown). As an amyloid is incessantly moving in a fine vibratory manner with twisting tendency, the hetero-oligomers might have discordance in the motions between the heterologous peptides that eventually lead to destabilization.

## Significance of behaviors of the stack-end molecules

MD simulations of various mutant αSyn amyloids demonstrated that behaviors of the stack-end molecules were highly varied depending on the primary structures, despite they were in the same conformation [52]. Whether the behaviors of the stack-end molecules (**Fig 2B, chain A or J**) affect incorporation of the substrate to the stack so as to pose a strain barrier is to be investigated but, if they do, they could modulate levels of barriers without substantial conformational changes depending on environments, e.g. pH or salt strength. Indeed, seeding reactions of amyloids are often affected by reaction conditions [38]. It can also explain why different strains have apparently similar secondary structures [62–64].

## Implications about prion/PrP^Sc^

We previously attempted to explain inter-species transmissions of prions from the view point that PrP^Sc^ is an in-register parallel β-sheet amyloid [54]. Indeed, in-register parallel β-sheet amyloids show many similar properties as PrP^Sc^. For example, the existence of different proteolytic-fragment patterns of Tau and αSyn amyloids [26][38][42] are reminiscent of the 19-kDa and 21-kDa PrP^Sc^ of prions [65]. Besides, those polymorphs of Tau or αSyn amyloids are associated with distinct clinicopathological pictures [10][42]. With the convenience in experiments, those proteins are good surrogate models for PrP^Sc^, which potentially provide clues to elucidation of the mechanisms of strain diversification and how strain-specific structures are translated into the manifestation.

The MD simulations of the mutant αSyn demonstrated that inappropriately high intrinsic β-sheet propensity of loops in certain shapes, e.g. U-shape, can destabilize the local structures of the amyloid. Inversely, local structures of a given amyloid might be inferable from influences of mutations around putative loop regions; if a mutation which raises local β-sheet propensity inhibits incorporation of the mutant peptide into the amyloid, the local structure around the mutation can be a U-shaped loop which cannot accommodate newly-induced β-sheets. Interestingly, the human PrP with an anti-prion polymorphism Val127 [66] shows positive (Pβ-Pc) values at the residues 126-127 where the PrP with Gly127 has negative values (Fig 4A). As the residue 127 is located at an end of the flexible region intervening two high-(Pβ-Pc) regions, -AVVGGLGG_127_YMLGS-, like the residue 84 of the αSyn amyloid, it may also affect structures of the loop and destabilize the surrounding structures. Moreover, as other residues associated with strain barriers are often located near presumably flexible regions, e.g., codon 129 of human PrP [65] and codon 136 of ovine PrP [67], the similar mechanism might affect the heterologous transmission efficiencies as well.

**Fig 4.**
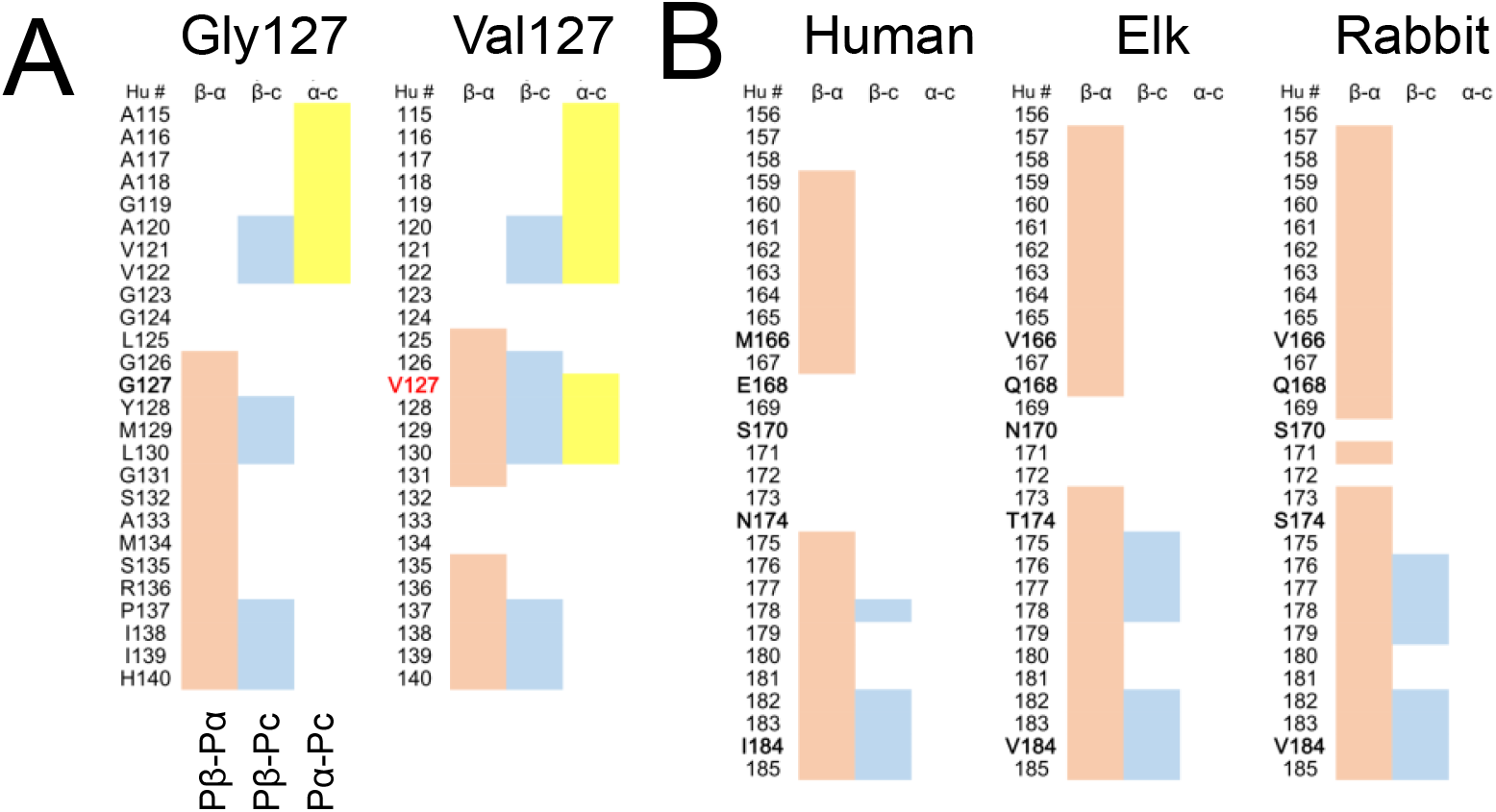
Influences of primary structures on the secondary structure prediction profiles of PrP. **A.** Comparison of the predicted propensities between two codon-127 polymorphs of human PrP, Gly127 (left) and Val127 (right). **B**. Comparison of the predicted propensities of H1∼H2 regions between human, elk and rabbit PrPs. Hu #, human numbering of the corresponding residues.

The region between the first and the second α-helices (H1∼H2) can be one of the main interfaces between substrate PrP^C^ and template PrP^Sc^, i.e., their first interaction site [68]. Although the predicted β-sheet propensities of this region is not necessarily high because of high coil propensity (**Fig 4B, left**), the conformation of native PrP^C^ possibly restricts the mobility of H1∼H2 region to make it more prone to β-sheet formation in effect than the predicted propensity. Remarkably, predicted β-sheet propensities of H1∼H2 are highly varied among species (**Fig 4B**), and the mismatches in the propensities in H1∼H2 may underlie species barriers by hampering stable β-sheet formation between PrP^Sc^ and PrP^C^ monomer; if the interface cannot convert to stable β-sheet structures, the conversion reaction would not spread to the rest of the molecule. Other regions with high β-sheet propensities, e.g. second or third α-helix, are stably structured in the native PrP^C^, and unsuitable for interfaces because they demand the energy to unfold before interaction with the template PrP^Sc^. There can be another interface in more N-terminal regions. Considering that Pro102Leu substitution of Gerstman-Sträussler-Scheinker syndrome (GSS) raises local β-sheet propensities in the N-terminal-side region, the 7-kDa protease-resistant fragment of GSS derived from the N-terminal region [69] may reflect preferential interactions of the region with PrP^Sc^ as a consequence of the raised local propensity. Interestingly, Fukuoka-1 mouse-adapted prion which was originally derived from GSS has distinct structures in H1∼H2 from that of other mouse-adapted strains RML or 22L [70].

## Implications about pathogenic mechanisms and conclusion

After strain-specific structures of amyloids are unveiled, the next question to be addressed is how those structures are translated into the strain-specific clinicopathological features. It is conceivable that the induction of conformational changes by the pathogenic amyloids also occur to contacting non-amyloidogenic proteins which has optimal intrinsic propensities, as if cross-seeding reactions between non-amyloidogenic protein and amyloid. If the same principles as that for between amyloidogenic proteins work, the non-amyloidogenic proteins which preferentially interact with an amyloid might be predictable from the primary structure.

Thus, we reviewed the recent advances in the investigations of detailed structures and mechanisms of strain diversity of in-register parallel β-sheet amyloids, particularly focusing on Tau and αSyn, and discussed their implications about prions/PrP^Sc^. Strain diversification of in-register parallel β-sheet amyloids seems to have multiple mechanisms due to intricate interactions between the β-sheets and due to local structures. As in silico approaches seem rather useful presumably because of the singular structural features, more proactive use of it would benefit the investigation.

## Acknowledgements

We thank Professor Kei Yura of Ochanomizu University for generously allowing us to use the neural network secondary structure prediction algorithm on his website. The numerical calculations were carried out on the TSUBAME2.5/3.0 supercomputer at the Tokyo Institute of Technology and the Reedbush-U supercomputer at the Information Technology Center, The University of Tokyo. This work was supported by “TSUBAME Encouragement Program for Young/Female Users” of Global Scientific Information and Computing Center at the Tokyo Institute of Technology, the “Initiative on Promotion of Supercomputing for Young or Women Researchers” from the Information Technology Center, The University of Tokyo, and Takeda Science Foundation (www.takeda-sci.or.jp/).

